# Cisplatin-inducible death receptor 5 antisense long non-coding RNA modulates cell cycle and proliferation in HeLa cells

**DOI:** 10.1101/2020.12.23.424136

**Authors:** Dilek Cansu Gürer, İpek Erdoğan, Ulvi Ahmadov, Osama Sweef, Bünyamin Akgül

**Affiliations:** Department of Molecular Biology and Genetics, İzmir Institute of Technology, Gülbahçeköyü, 35430 Urla Izmir, Turkey; Department of Biomedicine, Aarhus University, 8000 Aarhus, Denmark; Faculty of Science, Tanta University El-Gaish St., 31111 Tanta, Gharbia Governorate, Egypt

## Abstract

Cisplatin is a chemotherapeutic drug with pleiotropic effects known to modulate the expression of long non-coding RNAs (lncRNAs). With the annotation of many novel lncRNAs, it is imperative to employ a more comprehensive approach to identify cisplatin-mediated changes in the expression of lncRNAs. Next generation sequencing (NGS)-based profiling of total RNAs from cisplatin-treated HeLa cells identified 3489 expressed lncRNAs, of which 1930 and 1051 were up- and downregulated upon cisplatin treatment, respectively. For functional analyses, we selected one of the cisplatin inducible lncRNAs situated antisense to the death receptor 5 and thus named death receptor 5 antisense lncRNA (DR5-AS). Knock-down of DR5-AS lncRNA caused a morphological change in cell shape without inducing any cell death. A second round of NGS-based profiling of total RNAs from DR5-AS-silenced cells revealed differential expression of genes associated with the immune system and cell cycle. Further analyses showed that DR5-AS reduces cell proliferation and causes a cell cycle arrest at S and G2/M phases. These results suggest that cisplatin-mediated reduction in cell proliferation and cell cycle may be mediated by long non-coding RNAs.

**Significance:** Cisplatin is known to induce DNA-damage-induced cell death, which is used in combination chemotherapies in various cancer types. However, many patients develop resistance to cisplatin, which involves both protein-coding and noncoding genes. Although a number of long noncoding RNAs are linked to cisplatin resistance, a more comprehensive study is required. In this study, we took advantage of next-generation-sequencing based lncRNA profiling to unveil the extent of cisplatin inducible lncRNAs in HeLa cells. Additionally, we functionally characterized one of the cisplatin-inducible lncRNA, death receptor 5 antisense. Interestingly, this spesific lncRNA modulates cell morphology, proliferation and cell cycle without affecting cell death.

## Introduction

Cisplatin, a universal chemotherapeutic drug, is used in the treatment of a diverse array of cancer (1). As an alkylating-like agent, the platinum atom of cisplatin interacts with purines in DNA and induces crosslinks that lead to DNA damage and cell-cycle arrest (2). Such cellular perturbations trigger numerous signal transduction pathways and inflammatory pathways that trigger apoptosis. This DNA-damage-induced cell death is exploited in combination chemotherapies due to its synergistic effect. However, many patients develop resistance to chemotherapeutic drugs, including cisplatin (3). Thus, it is important to unravel the molecular mechanisms underlying the mode of action of platinum-based chemotherapeutic drugs.

Long non-coding RNAs (lncRNAs), non-protein-coding transcripts longer than 200 nt in length, are novel regulators of both transcriptional and post-transcriptional gene expression involved in numerous cellular processes such as growth, cell death and differentiation (4, 5). Transcribed largely by RNA polymerase II and III, several different biotypes of lncRNAs exist in the cell and they regulate gene expression by interacting with various macromolecules including DNA, RNA or proteins (6). The existing studies suggest that lncRNAs have the potential to modulate the molecular effect of cisplatin (7). Accordingly, bioinformatics analyses and microarray-based profiling of lncRNAs in different cancer cell lines and tissues treated with cisplatin clearly indicate that lncRNAs are involved in cisplatin-mediated cellular processes (8-10). Additionally, lncRNAs have been linked to cisplatin resistance through various molecular mechanisms that involve modulation of transcription (11), miRNA activity (12, 13), post-translational modification (14) or epigenetic silencing (15).

Despite the existence of several studies that associate lncRNAs with cisplatin’s mode of action, a robust profiling study is needed to obtain a comprehensive analysis of all types of lncRNAs. Here, we present an NGS-based lncRNA profile of HeLa cells treated with cisplatin. Our data show that cisplatin induces a broad repertoire of lncRNAs including lincRNAs, antisense lncRNAs and intronic lncRNAs. Death receptor 5 antisense (DR5-AS) lncRNA, a cisplatin-inducible natural antisense transcript (NAT) that is antisense to the DR5 receptor, modulates cell fate as its knockdown changes HeLa cell morphology. Transcriptomics analysis of DR5-AS-knocked-down cells has revealed that DR5-AS is involved in cell cycle and proliferation without affecting cell death.

## Results

### Cisplatin induces a plethora of long non-coding RNAs

Cisplatin, a chemotherapeutic drug with pleiotropic effects, is known to modulate several cellular properties. Thus, we first treated HeLa cells with varying concentrations of cisplatin to examine its effect on HeLa cells. Cisplatin at a concentration of 80 μM lowered the proliferation rate to 57.6% (Fig 1A) while inducing approximately 35.5% of apotosis, as indicated by Annexin V-positive early apoptotic cells, compared to the control DMSO (Figure 1B-D). To determine the extent of differentially expressed lncRNAs, we subjected three replicates of RNAs isolated from control and treated cells to RNA sequencing. The average read was 16,518,795. The heat map of differentially expressed transcripts is presented in Figure 2A. Cisplatin treatment resulted in the differential expression of 2981 lncRNAs (Figure 2B, 2-fold or higher, *P*<0.05). When we classified the differentially expressed lncRNAs based on their biotypes, we noticed that the majority of the differentially expressed lncRNAs were antisense lncRNAs and lincRNAs. However, it is quite interesting that many intron-derived lncRNAs were stably expressed upon cisplatin treatment. qPCR data of a select number of antisense lncRNA was congruous with the RNA-seq data (Figure 2C).

**Figure 1.**
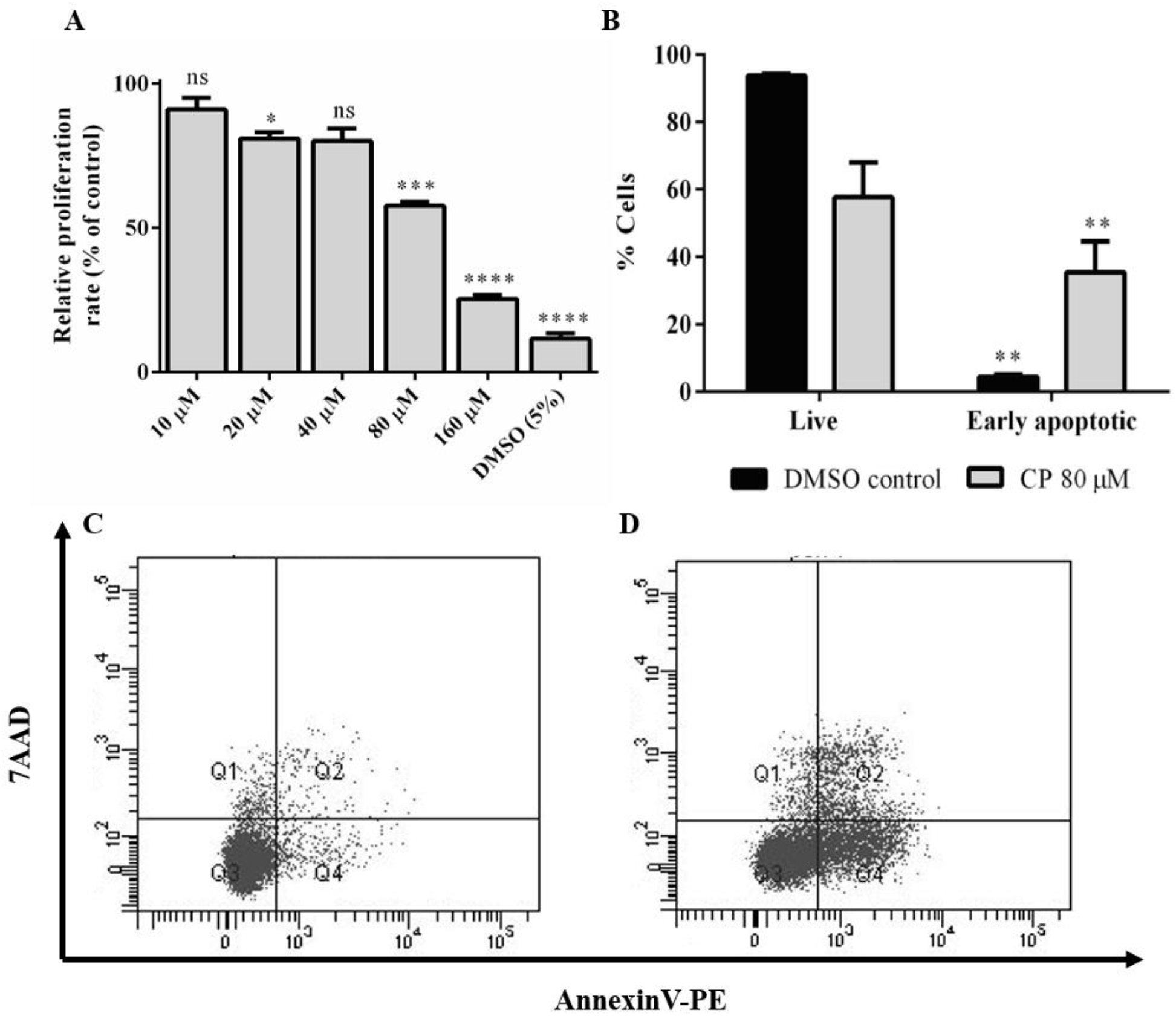
A. Proliferation kinetics of HeLa cells treated with a range of cisplatin concentrations. B. Percentage of live and early apoptotic population of HeLa cells treated with 80 μM cisplatin (CP). DMSO (0.1%) was used as negative control. C and D shows the population distributions of DMSO control and CP-treated groups as dot blot graphs, respectively. ns: non-significant, p>0.05, *: p≤0.05, **: p≤0.01, ***: p≤0.001, ****: p≤0.0001.

**Figure 2.**
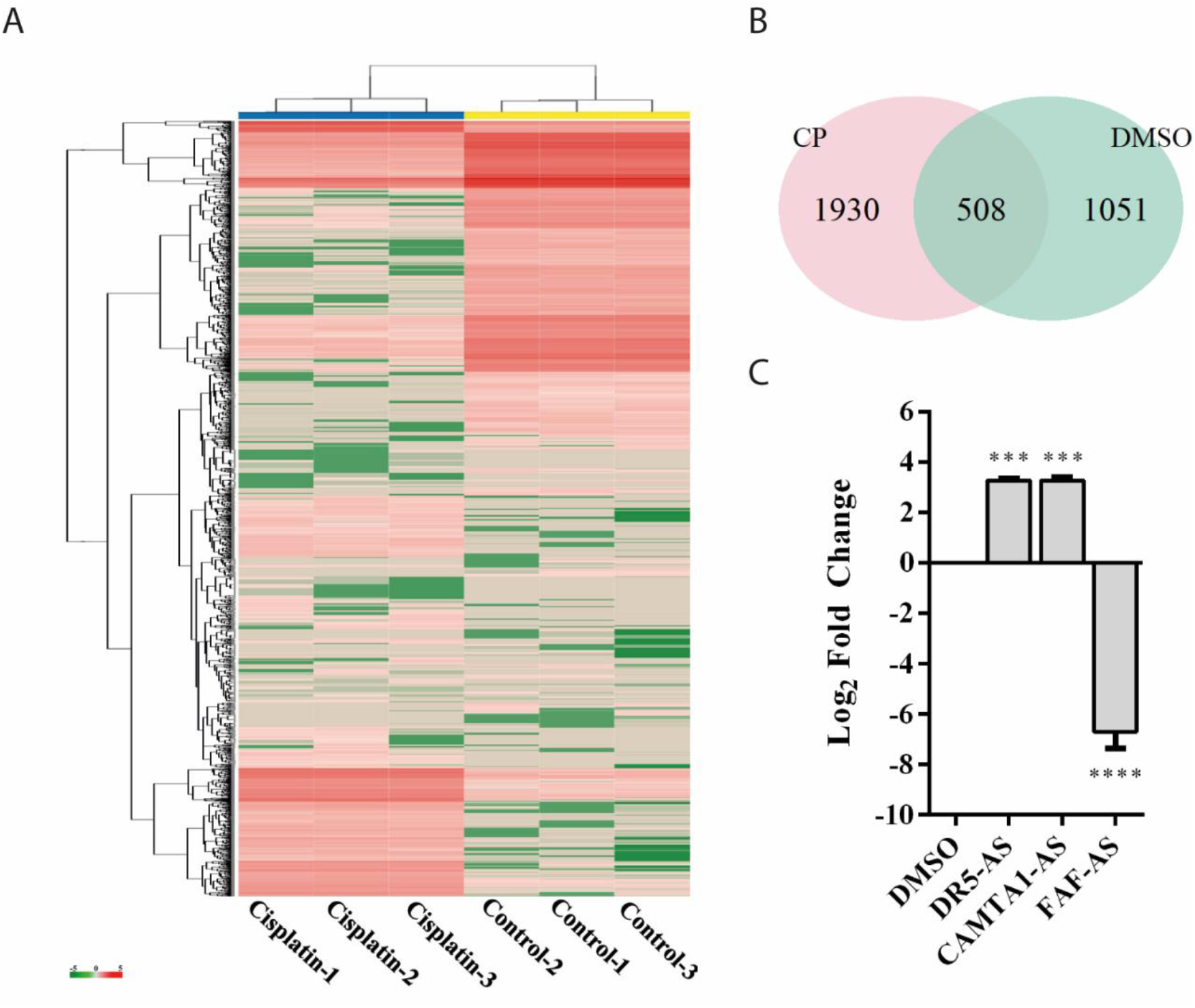
A. Heatmap of differentially expressed genes in cisplatin- and DMSO-treated HeLa cells. DMSO (0.1%) was used as control. B. Venn diagram indicating number of differentially expressed lncRNAs in cisplatin (CP) and DMSO-treated cells. C. qPCR analyses of candidate lncRNAs.

### DR5-AS is a cisplatin inducible nuclear antisense lncRNA

Of 2981 differentially expressed lncRNAs, DR5-AS stroke our attention because (1) it is antisense to the death receptor 5 (Figure 3A, 25); (2) it appears to be expressed in a tissue-specific manner (Supplementary Figure 1A); (3) mutations in this gene are associated with different types of cancer (Supplementary Figure 1B); (4) it appears to be fairly conserved (Supplementary Figure 1C); and (5) with 3 exons it is likely to be processed (Figure 3A). To examine the cell-specific expression pattern of DR5-AS lncRNA, we performed qPCR analyses with total RNAs isolated from two more cell lines, namely MCF7 and Jurkat cells. The expression of DR5-AS in HeLa and Jurkat cells were comparable but much less abundant in MCF7 cells (Figure 3B). Interestingly, cisplatin treatment induced the expression of DR5-AS lncRNA in HeLa and Jurkat cells but not in MCF7 cells (Figure 3C).

**Figure 3.**
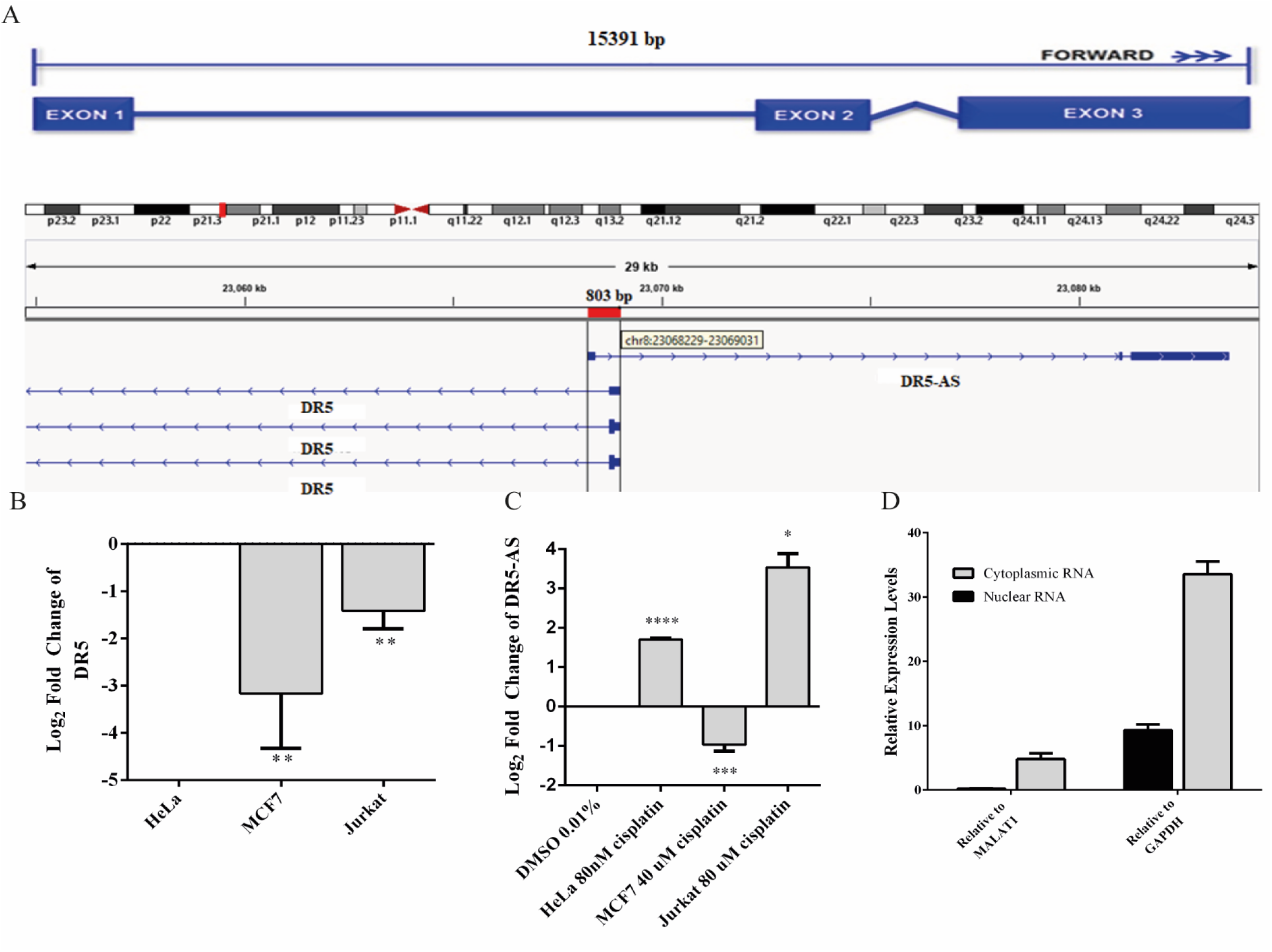
A. Schematic representation of DR5-AS and interference with DR5 transcript variant combined with the Ensemble structure. qPCR analysis of DR5-AS expression untreated (B and cisplatin-treated cells (C). D. Subcellular localization of DR5-AS. ns: non-significant, p>0.05, *: p≤0.05, **: p≤0.01, p***: p≤0.001.

The DR5-AS gene is annotated to encode a transcript with 3 exons (Figure 3A). To map the 5’ end 3’ borders of the DR5-AS transcript, we carried out 5’ and 3’ RACE. Our analysis showed that the DR5-AS gene encodes a 2636-nt transcript without a tail (Supplementary Figure 2). Since there appears to be a correlation between the subcellular location of an lncRNA and its regulatory function (26), we examined the intracellular distribution of this transcript. To this extent, qPCR analyses were performed with total, cytoplasmic and nuclear RNAs isolated from cisplatin-treated HeLa cells. To check for the integrity of subcellular RNA fractionation, we used MALAT1 and GAPDH as controls for nuclear and cytoplasmic RNA preparations, respectively (27). In reference to these markers, DR5-AS appears to be localized primarily in the nuclear fraction (Figure 3D).

### DR5-AS lncRNA modulates cell morphology

The DR5-AS lncRNA gene is physically overlapping with the protein-coding DR5 gene (Figure 3A). The DR5 receptor is known to trigger signal transduction pathways that modulate apoptosis, miRNA biogenesis, survival and proliferation (25). Considering the cisplatin inducibility of DR5-AS lncRNA and its genomic location antisense to the DR5 gene, we hypothesized that the DR5-AS lncRNA could be involved in modulating cell fate. To gain insight into the cellular function of DR5-AS, we exploited a reverse genetics approach to perturb intracellular DR5-AS lncRNA concentration. For this purpose, we employed the GapmeR technology as it has been reported to knock down nuclear lncRNAs more efficiently (28). We first quantified the intracellular amount of the DR5-AS lncRNA in HeLa cells 72h post-transfection with two different Gapmers (DR5-AS-GapmeR-1 and DR5-AS-GapmeR-2). qPCR analyses showed that GapmeR-1 was more efficient in knocking down the DR5-AS lncRNA (Figure 4A). We transfected cells with fluorescent-labeled DR5-AS-GapmeR-1 to examine the transfection efficiency. Fluorescent microscopy showed that more than 60% of HeLa cells were transfected under our transfection conditions (Data not shown). We also cloned the full-length cDNA of DR5-AS lncRNA into pcDNA3.1 to obtain pcDNA3.1-DR5-AS, which was efficiently overexpressed when transfected into HeLa cells (Figure 4A).

**Figure 4.**
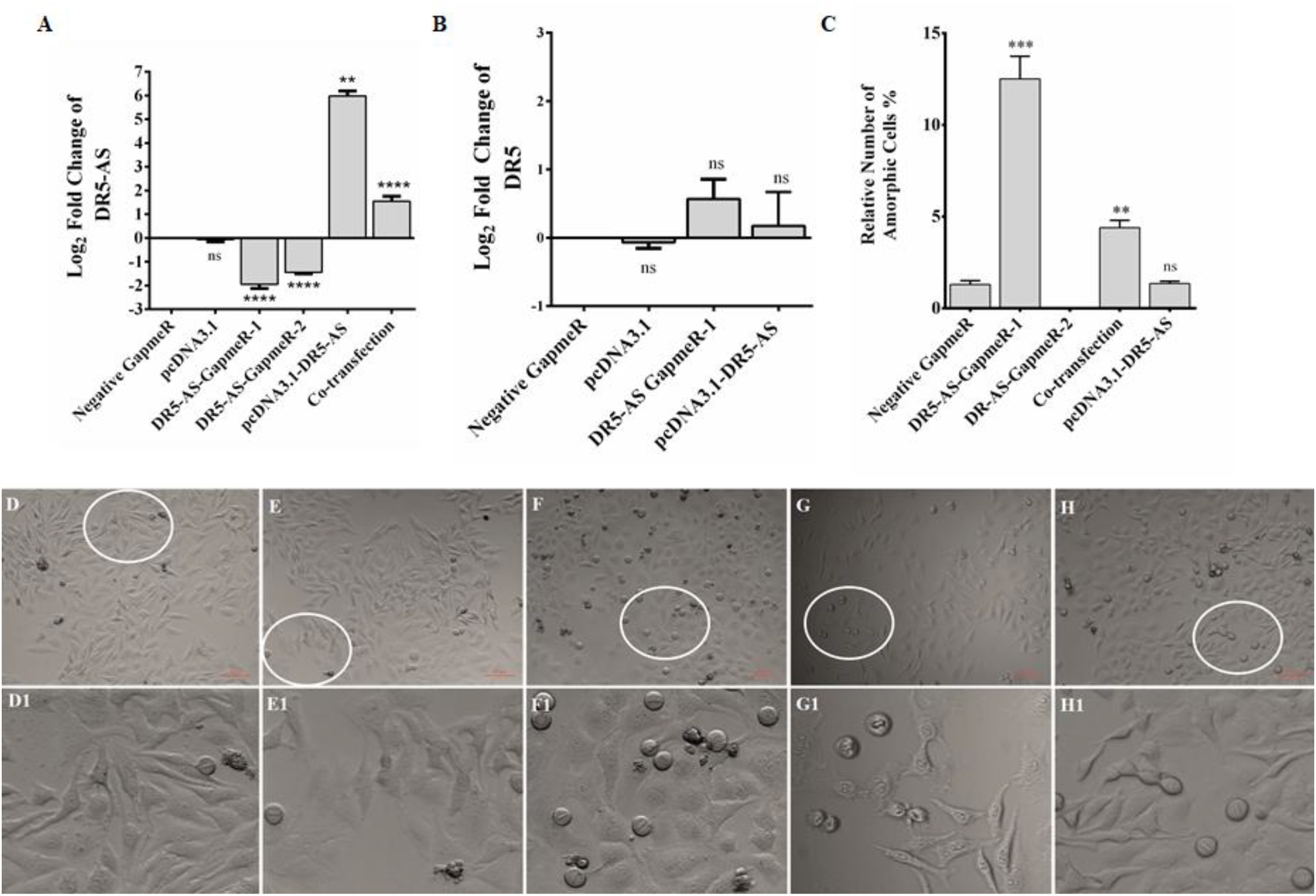
qPCR analyses DR5-AS (A) and DR5 sense mRNA (B) C. Quantitative number of metaphase block cells in DR5-AS silencing (DR5-AS-GapmeR), overexpression (pcDNA3.1-DR5-AS) and co-transfection groups. Negative GapmeR, represented by Neg-GapmeR was used as negative control for transfection. pcDNA3.1 represents the empty vector for overexpression. ns: non-significant, p>0.05, **: p≤0.01, ***: p≤0.001, ****: p≤0.0001. D-H. Brightfield images of transfected cells after incubation period. D1-H1 also represents the 300X magnified areas of the parts encircled in white. D and D1 indicates negative GapmeR, E and E1, pcDNA3.1-DR5-AS, F and F1, DR5-AS GapmeR-1, G and G1, DR5-AS GapmeR-2 and H and H1, co-transfection. Scale bar 50 μm.

A portion of antisense lncRNAs have been reported to regulate transcription of the protein coding gene with which it is overlapping (29). Based on the ENSEMBL entry, there is an 803-bp overlap between the DR5-AS lncRNA and the DR5 protein coding gene. To test the hypothesis that the DR5-AS transcript might transcriptionally regulate DR5 transcription *in cis*, we measured the amount of DR5 mRNA following DR5-AS knock-down and overexpression. Our qPCR analyses showed that DR5-AS knock-down or overexpression was not sufficient to perturb the intracellular DR5 mRNA amount under our experimental conditions (Figure 4B). Although we cannot conclusively eliminate the possibility that the native promoter-driven active transcription of DR5-AS may be required for regulation in *cis*, the mere increase in the DR5-AS transcript abundance does not appear to modulate the intracellular DR5 mRNA concentration. This finding is congruous with the earlier observation in that DR5-AS knockdown or overexpression does not influence the apoptotic rate of HeLa cells (Supplemental Figure 3A). Considering the fact that DR5 overexpression increases the rate of apoptosis in HeLa cells (30), a potential regulation in *cis* of DR5 by DR5-AS would be expected to ensue a change in the apoptotic rate.

We then monitored the cell morphology 72h post-transfection and observed a dramatic change in cell shape upon transfection of HeLa cells with 40 nM of two different versions of unlabeled GapmeR against DR5-AS (Figure 4C-H). Morphology of the cells transfected with negative GapmeR and pcDNA3.1-DR5-AS (Figure 4D and 4E, respectively) remained similar to characteristic adherent HeLa cells exhibiting projections from cytoplasm, while cells transfected with DR5-AS GapmeR-1 (Figure 4F), but not with DR5-AS GapmeR-2 (Figure 4G) were transformed into spherical-shaped adherent cells. These cells appeared to have been arrested at the metaphase as revealed by DAPI staining (Supplementary Figure 3). Approximately 12% of the cells possess this phenotype (Figure 4C). To ensure that the change in cell morphology is specifically due to the knock-down of DR5-AS, we also performed co-transfection assays with the DR5-AS overexpression construct, pcDNA3.1-DR5-AS. Although the overexpression of HeLa cells with pcDNA3.1.DR5-AS did not yield any observable cellular phenotype (Figure 4E), DR5-AS lncRNA overexpression was able to partially rescue the GapmeR-induced morphological change (Figure 4C and H). We then checked the viability of cells with morphological changes to eliminate the possibility that these cells are going through cell death. Our flow cytometric analyses showed that the percentage of Annexin V-positive cells were quite comparable in GapmeR-, pcDNA3.1.DR5-AS- and co-transfected cells, indicating that these cells are indeed viable (Supplemental Figure 4A). We also measured the intracellular uptake of NucRed™ Dead 647 ReadyProbes™ (ThermoFisher Scientific), a fluorescent dye used as a marker for dead cells. The NucRed™ Dead 647 ReadyProbes™ fluorescent dye penetrated into neither negative control GapmeR-nor DR5-AS-GapmeR-transfected HeLa cells, further confirming the viability of these cells (Supplemental Figure 4B).

### DR5-AS knock-down perturbs the transcriptome associated with cell proliferation and cell cycle

Cisplatin is known to exert a pleiotropic effect on cells (31). Based on the physical overlap between DR5-AS and DR5 with respect to their genomic location, any functional interaction would be expected to manifest a change in the apoptosis rate. Our inability to detect such a phenotypic outcome (Supplemental Figure 4) suggests that DR5-AS may function through other cellular processes, such as stress, proliferation, drug metabolism or cell motility, all of which are known to be modulated by cisplatin treatment.

To gain insight into the cellular function of the DR5-AS transcript, we exploited the power of the transcriptomics approach. To this end, we first knocked-down the DR5-AS transcript with GapmeR-1 and sequenced the total RNAs isolated from these cells in parallel to those isolated from control-Gapmer-transfected HeLa cells. Bioinformatics analyses revealed the differential expression of 2215 mRNAs, of which 876 and 1339 up- and down-regulated, respectively (Figure 5A-B). We then performed a Reactome pathway analysis to deduce biological processes affected by DR5-AS knockdown. Strikingly, this analysis showed that DR5-AS is very likely to be involved in immune system-related cellular processes (Figure 5C). Interestingly, we have identified two more biological processes, cell cycle and proliferation, that could be pertinent to the cisplatin inducibility of the DR5-AS transcript (Figure 5C). Thus, we validated by qPCR the amount of some DEGs associated with cell proliferation and cell cycle. The qPCR results were in congruous with the RNA-seq data except for C5 (Figure 5D).

**Figure 5.**
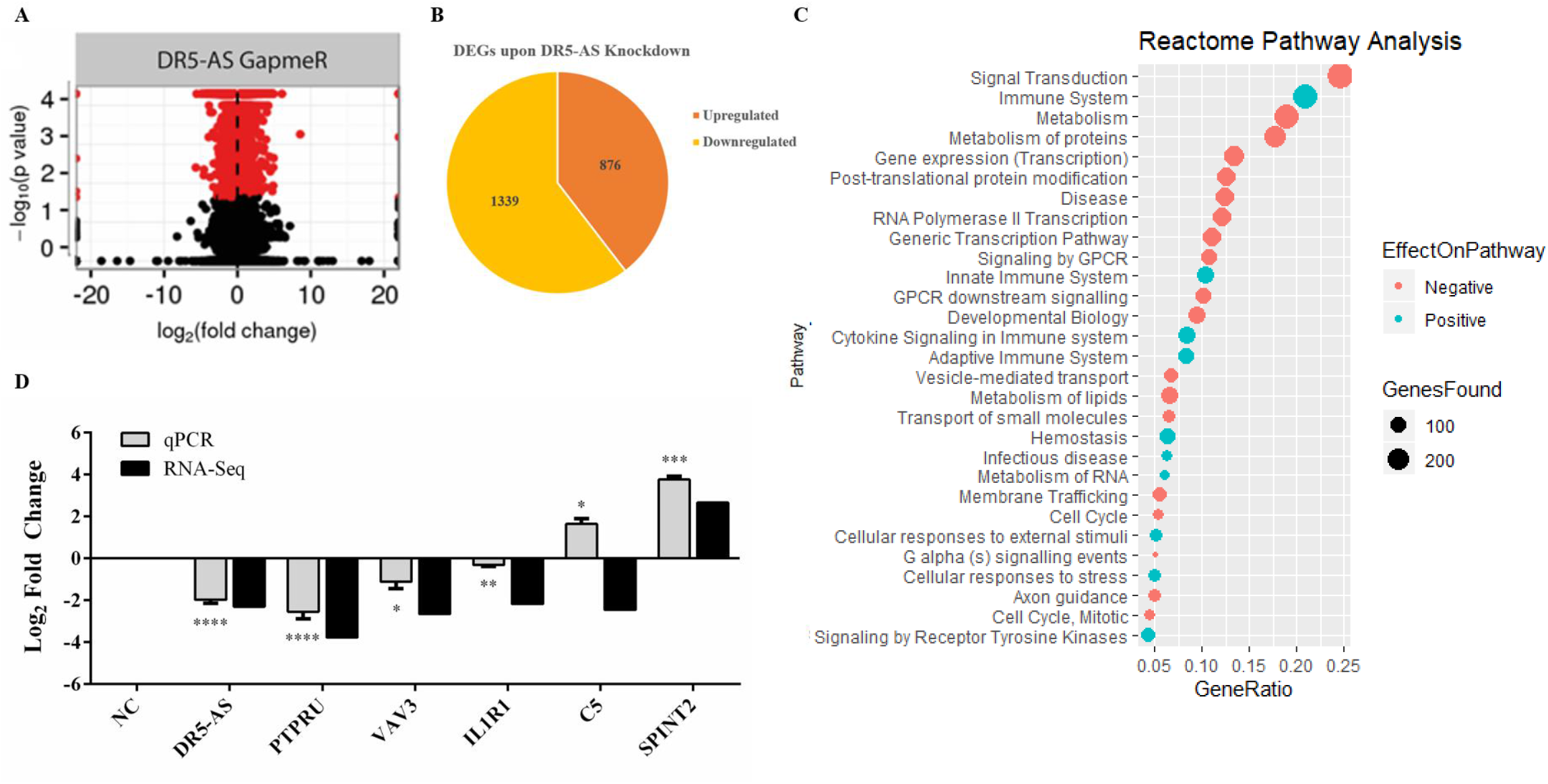
A. Volcano plot and B. Pie chart of differentially expressed genes (DEGs) after knock-down. C. Pathway enrichment analysis of DR5-AS knockdown RNA-Seq data by using Reactome Pathway Database. D. qPCR validation of candidate genes. *: p≤0.05, **: p≤0.01, ***: p≤0.001, ****: p≤0.0001.

To functionally test the observations made by the transcriptomics approach, we measured the proliferation rate of HeLa cells following 72h transfection with negative and DR5-AS GapmeR. In agreement with the transcriptomics data, we observed a 22.7% reduction in the proliferation rate of HeLa cells upon DR5-AS knock-down in comparison with the cells transfected with negative GapmeR (Figure 6A). There was a correlation between the knock-down efficiency of GapmeR-1 and 2 (Figure 4A) and the corresponding reduced proliferation rate. To ensure that the GapmeR-mediated knockdown was responsible for this reduction in proliferation rate, we tried to rescue the phenotype by overexpressing DR5-AS lncRNA. As expected, the overexpression of DR5-AS has partially rescued the DR5-AS-GapmeR-mediated decrease in the proliferation rate (Figure 6A).

**Figure 6.**
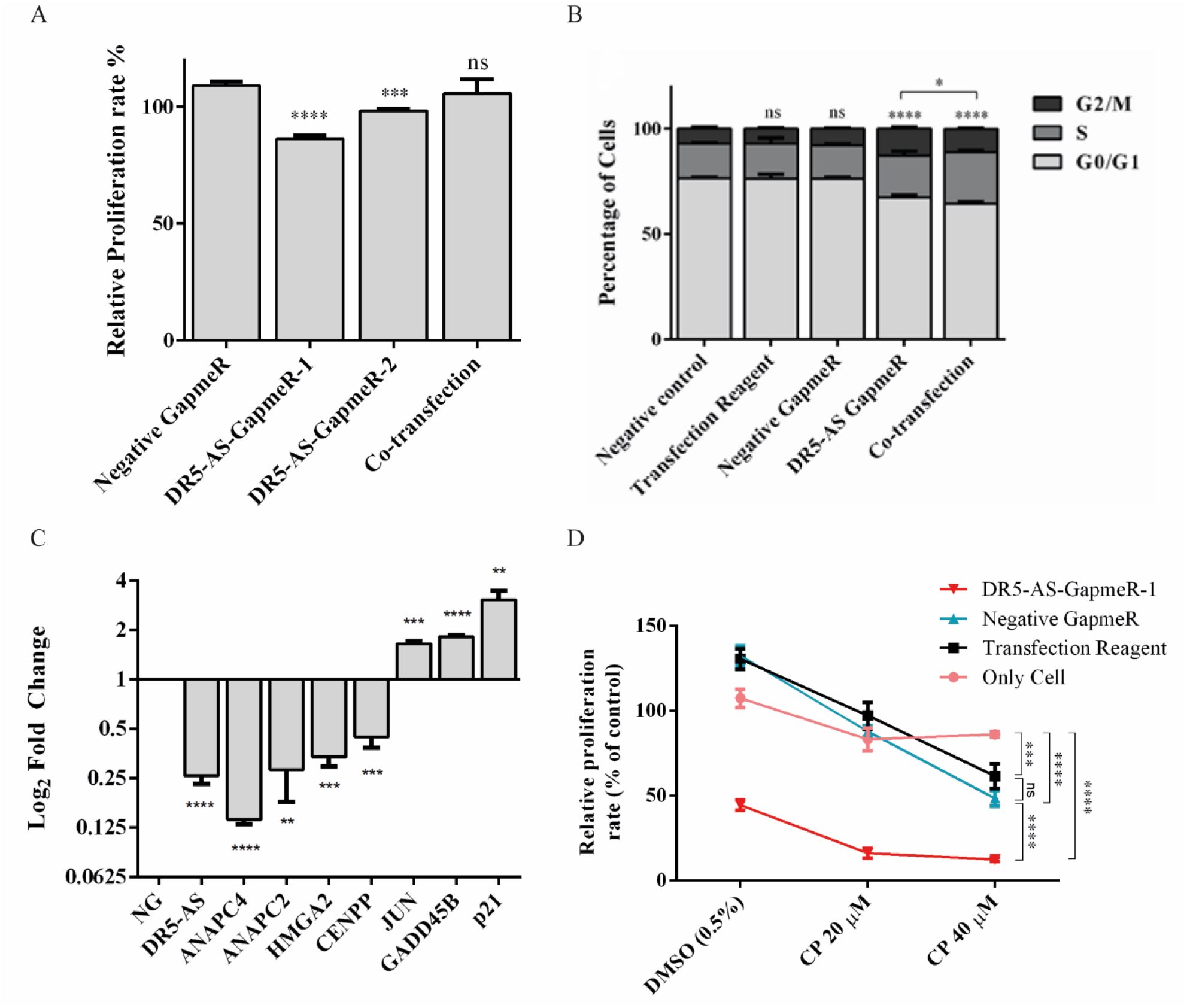
A. Proliferation and B. Cell cycle analysis after silencing and co-transfection. C. qPCR analyses of genes associated with cell cycle and/or proliferation. D. Proliferation rate of cells after cisplatin treatment followed by DR5-AS knock down. Statistical analyses were performed by student’s t-test for A,B,C,D and two-way ANOVA with Tukey’s multiple comparison for D. ns: non-significant, p>0.05, *: p≤0.05, **: p≤0.01, ***: p≤0.001, ****: p≤0.0001.

We extended our functional analysis to cover cell cycle analyses as well since DR5-AS knock-down perturbed the gene expression pattern associated with cell cycle (Figure 5C). Thus, we performed a post-knock-down flow cytometric cell cycle analysis. Although the transfection reagent or the negative GapmeR did not cause any discernible difference in the cell cycle profile, DR5-AS knock-down caused the cells to shift from the G0/G1 phase to the S and G2/M phases (Figure 6B). Although DR5-AS overexpression did not completely rescue the DR5-AS-knock-down-mediated shift, it has reduced the percentage of the cells in the G2/M phase from 12.7% to 11%. We then selected a number of DEGs associated with cell cycle that were identified through the reactome pathway analyses (Figure 5C) for validation by qPCR. Parallel to RNA-seq data, DR5-AS knock-down induced p21 and GADD45B and down-regulated the expression of ANAPC2 and ANAPC4 which are subunits of anaphase promoting complex/cyclosome (APC/C) and CENPP, as key regulators of cell cycle (Figure 6C) (32). To test whether DR5-AS knock-down exacerbates the antiproliferative effect of cisplatin, we first transfected HeLa cells with the DR5-AS and then treated with relatively milder concentrations of cisplatin (e.g., 20 and 40 μM). DR5-AS knock-down exacerbated the antiproliferative effect of cisplatin nearly 3.2-fold (Figure 6D).

## Discussion

We provide the first comprehensive expression profile of lncRNAs in cisplatin-treated HeLa cells. Under our experimental setting, we identified 2981 de-regulated lncRNAs that include not only the well-known examples of lincRNAs and antisense ncRNAs but also an interesting repertoire of intronic lncRNAs. We selected DR5-AS for further functional analysis because of its genomic location being antisense to the death receptor (DR) 5, a TRAIL-bound receptor that modulates cell death and survival (33). GapmeR-mediated silencing of the nuclear DR5-AS transcript caused a change in cell morphology without affecting cell viability, which could be partially rescued upon its ectopic overexpression. Transcriptomics analyses of the DR5-AS-knocked-down HeLa cells uncovered changes in the expression of genes associated with cell morphology, cell proliferation and cell cycle. Congruently, DR5-AS knock-down led to a drop in the proliferation rate and brought about a shift in the cell cycle as evidenced by an arrest at S and G2/M phases.

Previous microarray studies have shown that cisplatin modulates the expression of lncRNAs in various cisplatin-treated cell lines such as A549 lung adenocarcinoma cell line (8, 9) and CAL-27 and SSC-9 tongue squamous carcinoma cell lines (11). In addition, bioinformatics methods were employed to examine lncRNAs associated with platinum drugs in high grade serous ovarian cancer (34). Interestingly, a pan-cancer analysis of RNA-seq data of 648 samples from 11 different cancer types revealed cancer-specific lncRNA expression upon cisplatin treatment. Subsequent functional analyses have shown that lncRNAs could function through various mechanisms to modulate cisplatin-modulated cellular responses. For example, cisplatin-sensitivity-associated lncRNA (CISAL) inhibits BRCA1 transcription and thereby controls cisplatin sensitivity in squamous cell carcinoma (11). LncRNA AK126698 was reported to be involved in Wnt/B-catenin-mediated regulation of cisplatin-induced apoptosis in A549 cells (8). LncRNA TUG1, on the other hand, promotes cisplatin resistance through the epigenetic regulation of miR-194-5p in bladder cancer (15). Although microarray-based transcriptomics approaches have been highly fruitful in identification of especially highly expressed lncRNAs, they are limited to the analysis of known lncRNAs. We employed an NGS-based approach to ensure a more comprehensive coverage. Indeed, our analyses revealed the stable accumulation of a number of intron-derived lncRNAs.

Antisense lncRNAs have been reported to regulate the expression of nearby genes in *cis* or *trans* (29). In addition to the regulation of transcription-related processes, NATs can modulate gene expression through RNA:DNA or RNA:RNA interactions in the nucleus or RNA:RNA interactions in the cytosol (35). We prioritized DR5-AS for functional analyses as it is positioned antisense to the DR5 receptor. TNF-related apoptosis inducing ligand (TRAIL), which is an anticancer agent in human, triggers apoptosis by ligating to the DR5 receptor (36), making DR5-AS a prominent choice for studying cisplatin’s downstream effects. Although DR5-AS is located in the nucleus (Figure 3), neither its ectopic overexpression nor its GapmeR-mediated knockdown modulates DR5 expression under our experimental setting (Figure 4B). This suggests that either DR5-AS regulates its target genes in *trans* or its in-*cis* transcription is required to regulate the sense mRNA.

We exploited reverse genetics to gain insight into the potential function of DR5-AS. To this extent, we employed GapmeR technology as GapmeRs are more efficient in knocking down nuclear lncRNAs compared to siRNAs (28). Interestingly, knocking down DR5-AS caused a severe change in cell morphology in HeLa cells (Figure 4F-F1), which could be partially rescued by overexpression of DR5-AS (Figure 4H-H1). These round-shaped cells maintained adherence to the culture flasks and were alive as evident by flow cytometric analysis of cells (Supplementary Figure 4A) and by microscopic observation after NucRed™ Dead 647 ReadyProbes™ staining (Supplementary Figure 4B). Surprisingly, RNA-seq analysis of total RNAs from DR5-AS-knocked-down cells revealed modulation of immune-system-related genes (Figure 5C) in HeLa cells of epithelial morphology. Considering the GapmeR-mediated morphological change and relevance to cisplatin’s effects, we noticed the changes in gene expression associated with cell cycle and proliferation. Expectedly, DR5-AS knock-down resulted in a decrease in the proliferation rate of HeLa cells coupled with a cell cycle arrest (Figure 6). It is highly interesting that DR5-AS knock-down reduces proliferation rate and causes a cell cycle arrest without triggering cell death. This phenomenon is typically seen in the immune system. For example, increased cell proliferation, without a change in cell death, has been reported to induce lymphocytosis in bovine leukemia virus-infected sheep (37). Taking into account the extent of affected immune system-related genes following DR5-AS knock-down, it would be interesting to probe into the potential function of DR5-AS in regulating innate or adaptive immunity.

## Materials and Methods

### Cell Culture and Drug Treatments

HeLa cells, obtained from DKFZ GmbH (Germany), were cultured in RPMI 1640 (with 2 mM L-Glutamine, Gibco, USA) supplemented with 10% fetal bovine serum (FBS) (Gibco, USA) and 100 U/ml penicillin in combination with 100 μg/ml streptomycin (Gibco) in a humidified atmosphere of 5% CO_2_ at 37°C. After optimization of dose and time kinetics for apoptosis rate, cisplatin treatments were carried out in triplicates as described previously (16).

### Total RNA Isolation, RNA-Seq and qPCR

Total RNA was isolated using TRIzol (Life Technologies) according to the manufacturer’s instructions. Nuclear and cytoplasmic RNAs were isolated using the cytoplasmic and nuclear RNA purification kit (Norgen Biotek, Canada). Trace DNA contamination was removed with the TURBO DNA-free™ kit (Invitrogen, United States).

Five μg total RNA was used for library preparation and run on Illumina HiSeq2500 by FASTERIS to identify differentially expressed lncRNAs (https://www.fasteris.com/dna/). The RNA-seq data were analyzed by the OmicsBOX bioinformatic platform in which Spliced Transcripts Alignment to a Reference (STAR) aligner was used for the genome-guided alignment during transcript reconstruction (17). DEseq2 was employed to identify differentially expressed transcripts (18). Gene ontology analyses were carried out using the PANTHER program (19).

To identify differentially expressed mRNAs in DR5-AS-silenced HeLa cells, a similar RNA sequencing was conducted with total RNAs (3 replicates) isolated from control and DR-5-AS-silenced cells (Fasteris SA). RNA-seq data were deposited into the Gene Expression Omnibus under the accession number GSE160227 Differentially expressed mRNAs were subjected to Pathway Enrichment Analysis by Reactome database (20) and visualized with ggplot2 package in R platform (21).

For qPCR analyses, cDNA was prepared using RT^2^ first strand kit according to manufacturer’s instructions (Qiagen, United States). qPCR reactions were prepared with RT^2^ SYBR master mixes and RT^2^lncRNA qPCR assays for DR5-AS (Qiagen Cat., LPH15855A-200), CAMTA-AS (Qiagen Cat., LPH13091A-200) and FAF1-AS (Qiagen Cat., LPH05521A-200). GAPDH (Qiagen Cat., LPH31725A-200) was used for normalization.

### RACE and Construction of Overexpression Vectors

RACE-cDNA was synthesized according to the manufacturer’s protocol for 5’/3’ RACE Kit 2^nd^ generation (Roche; Cat.No. 03353621001). The 2648-bp cDNA was cloned into the *NheI-XhoI* site in pcDNA 3.1 plasmid to obtain pcDNA3.1-DR5-AS and was verified by sequencing. The empty vector was used as the negative control. Amplified plasmids were isolated using an endotoxin-free plasmid isolation kit (Macherey-Nagel, Germany).

### Cell Transfection

Cells were seeded in 6-well plates at a density of 80,000 cells/well one day prior to transfection and transfected with DR5-AS LNA GapmeR (Qiagen, United States) at a concentration of 40 nM or with 1500 ng of pcDNA3.1-DR5-AS with the Fugene HD transfection reagent (Promega, United States) in a 2-mL final volume. Negative GapmeR and empty vector were used as negative control groups for silencing and overexpression, respectively. Media was changed one hour post-transfection in overexpression experiments. Overexpression for rescue experiments was conducted 8h post-silencing under conditions similar to individual transfections. Transfected cells were fixed with methanol for brightfield microscopy and stained with DAPI for fluorescent microscopy (22). Cell morphology was analyzed with Fiji (23).

### Analysis of Apoptosis, Proliferation and Cell Cycle

To determine apoptotic cells, biological replicates were trypsinized by 1X Trypsin-EDTA (Gibco, United States) and washed in 1X cold PBS (Gibco, United States), followed by resuspension in 1X Annexin binding buffer (Becton Dickinson, United States). The resuspended cells were stained with Annexin V-PE (Becton Dickinson, United States) and 7AAD (Becton Dickinson, United States) followed by incubation for 15 minutes in dark at room temperature and analyzed by FACSCanto (Becton Dickinson, United States). Additionally, NucRed™ Dead 647 ReadyProbes™ (Invitrogen, United States) were used to assess the viability of the transfected cells through a fluorescent microscope (Zeiss Observer Z1, Germany). To this end, 2 drops of the dye were applied per mL of culture medium and cells were monitored following 30 min of incubation in dark.

WST-1 assay was employed to measure proliferation rates (Roche, Switzerland). Cells were seeded at a density of 1000 cells per well in 96-well plates one day prior to transfection. Transfection was carried out when the confluency has reached approximately 70-80% in a volume of 100 μl per well. Samples were incubated in a humidified atmosphere of 5% CO_2_ at 37°C for 2 hours following the addition of WST-1 and absorbance was measured at 450 nm. DMSO (5%) (v/v) was used as positive control while media without cells was used as blank.

For cell cycle analysis, trypsinized cell pellets were fixed with cold ethanol, permeabilized with Triton X-100 (0.1%) and treated with RNase A (24). The resulting cells were then stained with propidium iodide (PI) (Becton Dickinson, United States) for 15 min prior to analysis with FACSCanto (Becton Dickinson, United States). The population density in each cell cycle phase was calculated by ModFit LT software.

## Acknowledgements

This study was funded by TUBITAK (Project No: 117Z243 to BA). The authors would like to thank Devrim Pesen Okvur for technical help, Özgür Okur and Dane Ruscuklu for flow cytometry analyses and sequencing, and BIOMER (IZTECH, Turkey) for the instrumental help.

---

**Supplementary Figure 1.**
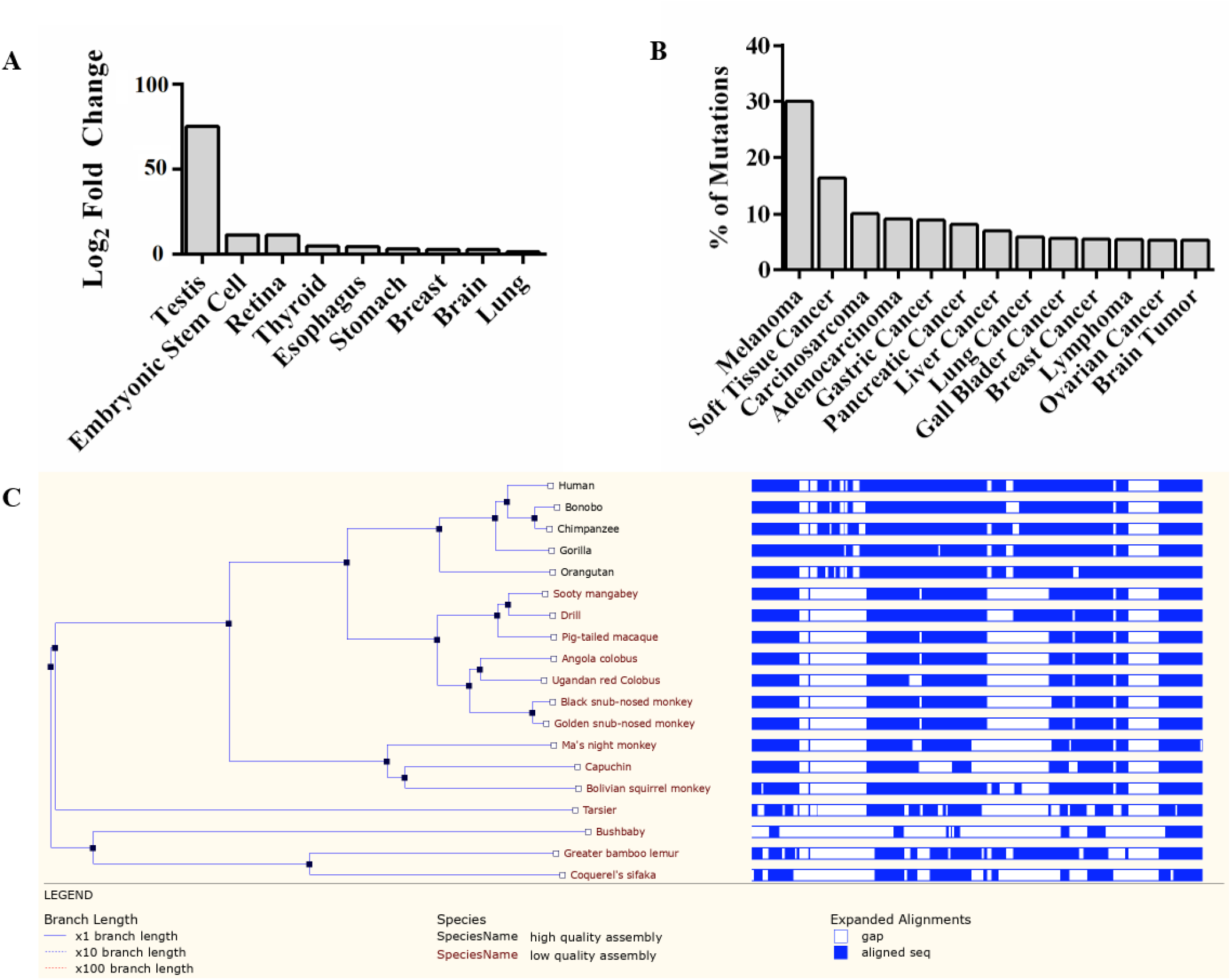
A. Differential expression and B. Percentage of mutations observed in DR5-AS in different cancer cases. C. MSA analysis of DR5-AS by 27 primates EPO-Low-Coverage.

**Supplementary Figure 2.**
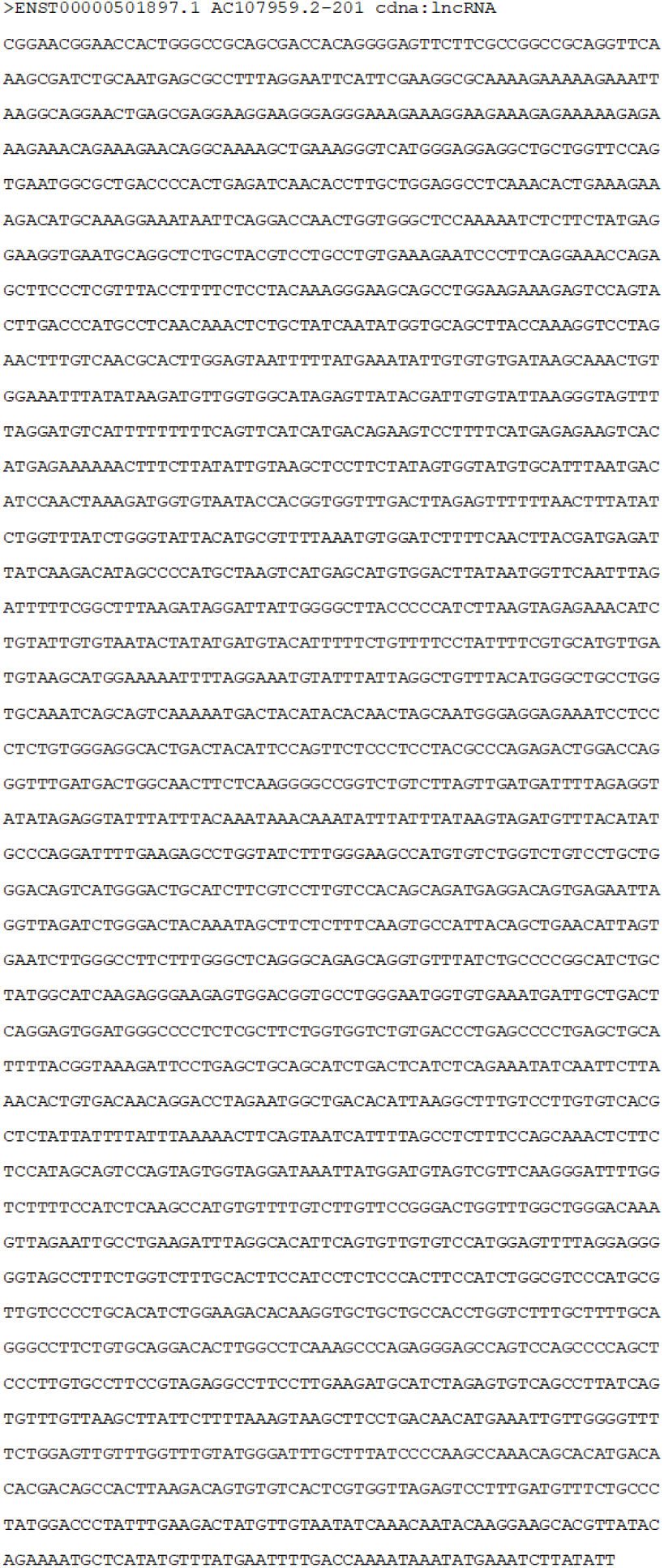
Full-length cDNA sequence of DR5-AS.

**Supplementary Figure 3.**
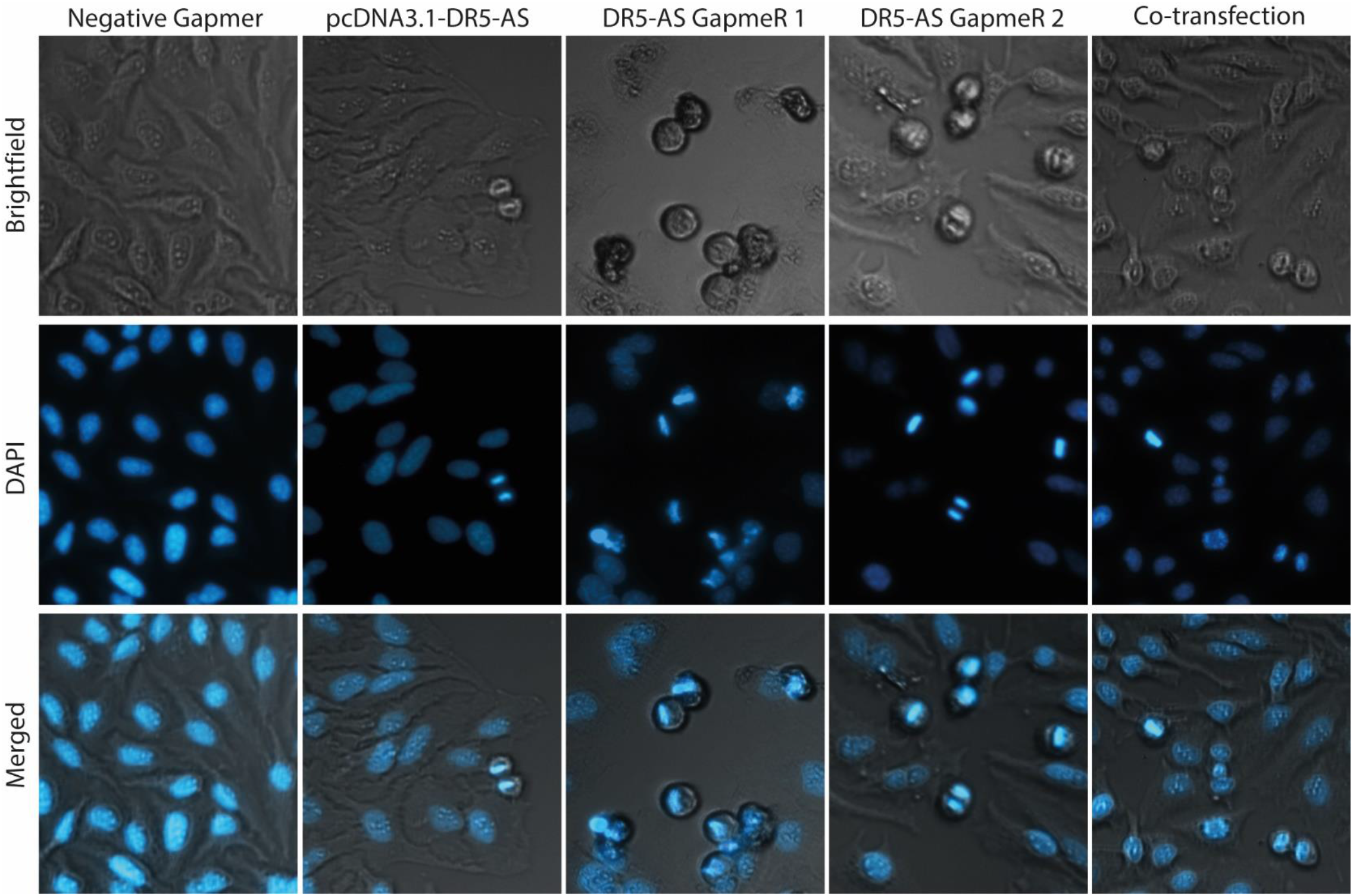
Phenotypic observation of DR5-AS-silenced HeLa cells stained with DAPI. Negative Gapmer was used as a negative control. Magnification is 10X

**Supplementary Figure 4.**
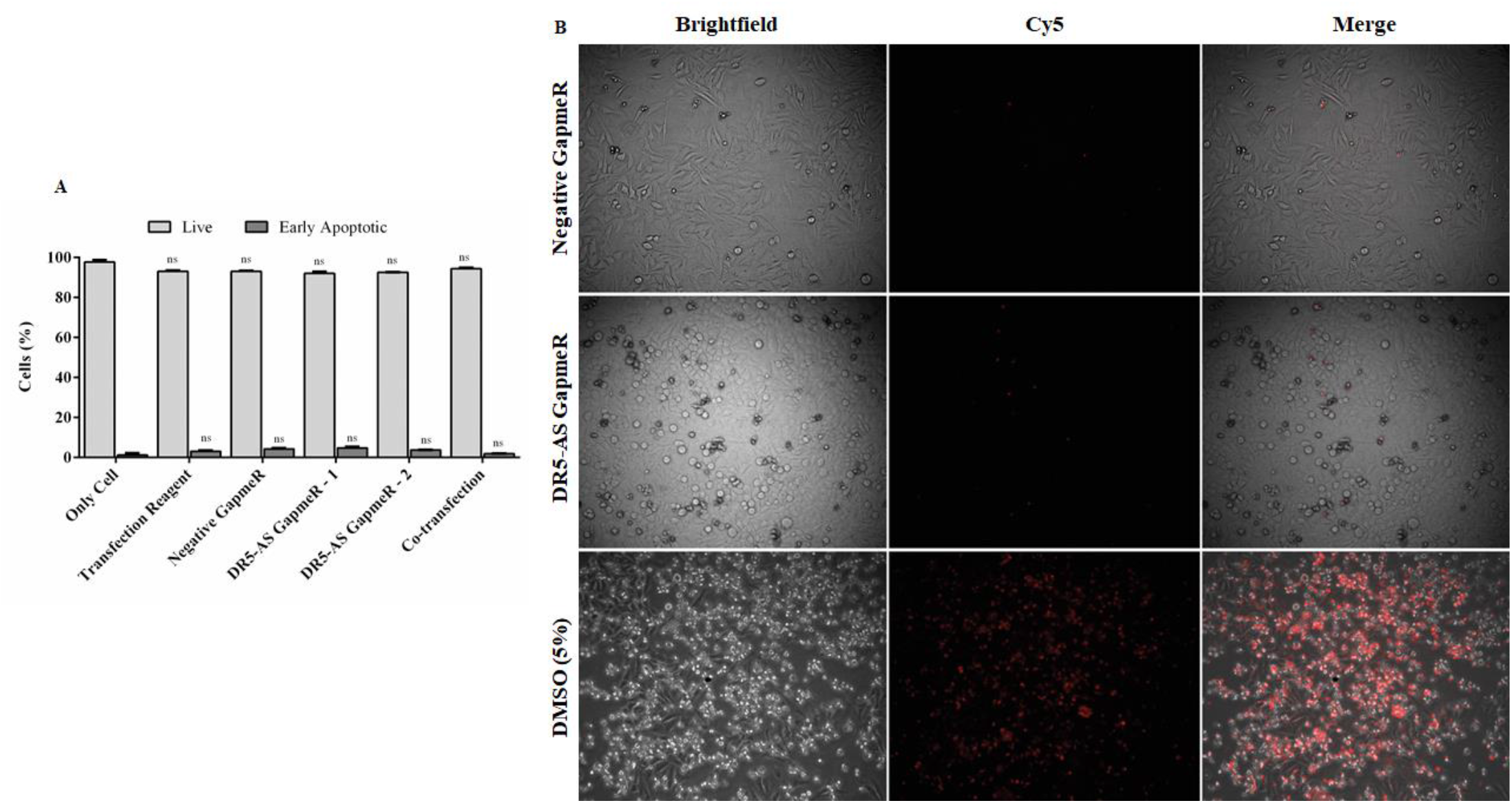
Flow cytometric and phenotypic observation after GapmeR-assisted silencing. A. Percentage of live and early apoptotic population. ns: non-significant, p>0.05. B. Live cell imaging by NucRed™ Dead 647 ReadyProbes™. DMSO (5%) was used as positive control. Magnification is 10X.

**Table 1:**
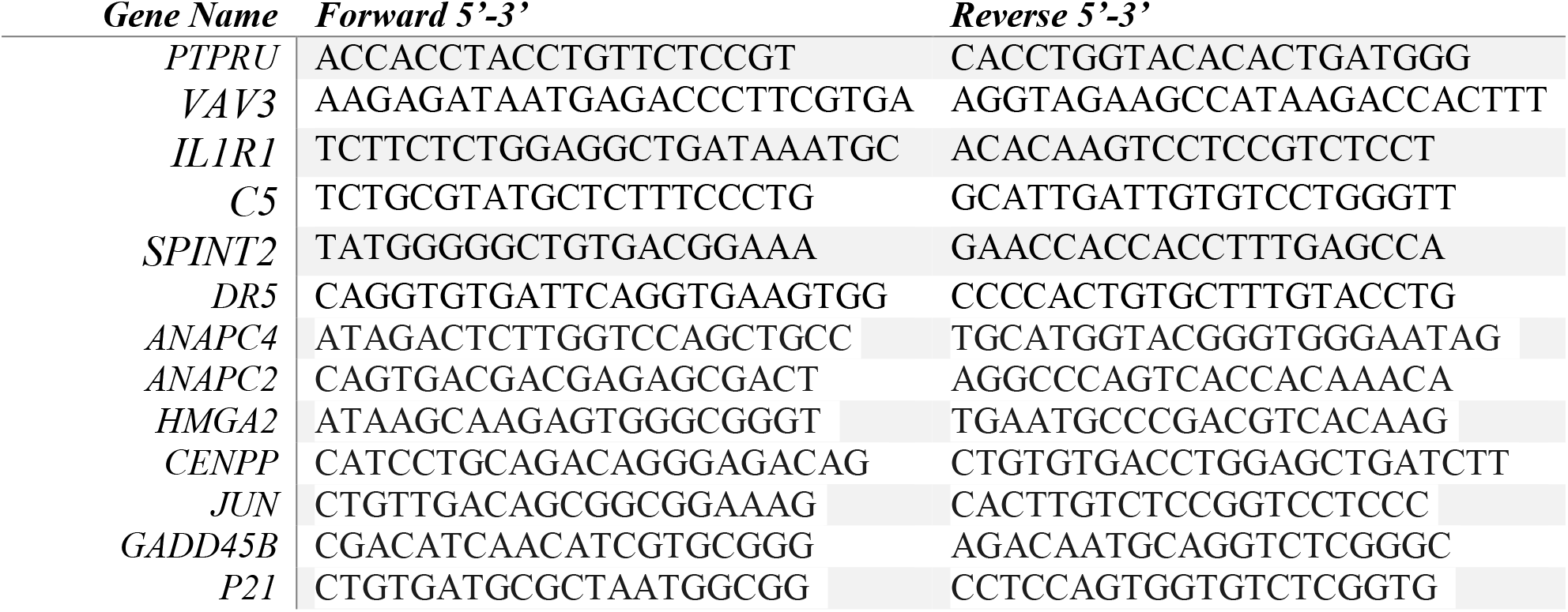

